# A Comprehensive Integrated Anatomical And Molecular Atlas Of Rodent Intrinsic Cardiac Nervous System

**DOI:** 10.1101/661033

**Authors:** Sirisha Achanta, Jonathan Gorky, Clara Leung, Alison Moss, Shaina Robbins, Leonard Eisenman, Jin Chen, Susan Tappan, Maci Heal, Navid Farahani, Todd Huffman, Steve England, Zixi (Jack) Cheng, Rajanikanth Vadigepalli, James S. Schwaber

## Abstract

In this study, we developed, coordinated, and integrated several technologies including novel whole organ imaging, software development to support the very first precise 3D neuroanatomical mapping and molecular phenotyping of the intracardiac nervous system (ICN). While qualitative and gross anatomical descriptions of the anatomy of the ICN have been presented, we here bring forth the first comprehensive atlas at large scale of the entire ICN in rat at a single cell resolution. Our work <u>*for the first time*</u> provides a novel 3D model to precisely integrate anatomical, functional and molecular data in the 3D digitally reconstructed whole heart with high resolution at the micron scale. This work represents the cutting edge in a long history of attempts to understand the anatomical substrate upon which the neuronal control of cardiac function is built. To our knowledge, there has not yet been a comprehensive histological mapping to generate a neurocardiac atlas at cellular and molecular level for the whole heart of any species. We now display the full extent and the position of neuronal clusters on the base and posterior left atrium, and the distribution of molecular phenotypes in that context. In addition we display in this context distinct molecular phenotypes that are defined along the base-to-apex axis, and the present novel discovery of their phenotypical spatial gradients, have not been previously described. The development of these approaches needed to acquire these data has produced method pipelines which can *not only* achieve the goals of anatomical and molecular mapping of the heart, but also provide the method pipelines for mapping other organs (e.g., stomach, lung, kidney, and liver).

## INTRODUCTION

In recent years neuroanatomy research in mammalian brain has come to the forefront, (e.g. in the BRAIN Project, Blue Brain Project and Connectome Project) dependent on the development of three-dimensional (3D) digital reference atlases at cellular scale, and the revelation of the complex diversity of the molecular phenotypes of neurons, often showing orderly spatial gradients of neuron types (e.g. work from the David Van Essen lab and the Allen Institute). In this report we further develop and extend these approaches to bring them to bear on the rodent intrinsic cardiac nervous system (ICN). Here, we present a comprehensive 3D mapping of rat ICN distribution in the overall histological and ontological context of the heart while demonstrating anatomically specific single-neuron transcriptional gradients. We herein show the development, coordination, and integration of several technologies including novel whole organ imaging, software development, precise 3D neuroanatomical mapping and molecular phenotyping. The development of the approaches needed to acquire these data has produced two method pipelines that can achieve the goals of the National Institutes of Health Common Fund’s Stimulating Peripheral Activity to Relieve Conditions (SPARC) program: Comprehensive Functional Mapping of Neuroanatomy of the Heart.

Autonomic control of cardiac function arises from several integrative centers in the central nervous system, but the final level of neural integration controlling cardiac function lies in the intrinsic cardiac nervous system (ICN). While the heart is known to possess a significant population of neurons, these have not previously been mapped precisely as to their number, extent/position/distribution while maintaining the histological context of the whole heart. This 3D neuroanatomical information is necessary to understand the connectivity of the neurons of the ICN and to develop their functional circuit organization. Mapping of molecular phenotypes and cell functions must also occur in the anatomical context to ensure a holistic comprehension of ICN.

Prior mapping efforts of ICN in small animals have included: mouse (Li et al., 2010, 2014; Rysevaite et al., 2011a), rat (Ai et al., 2007; Cheng et al., 1999, 2004; Cheng and Powley, 2000), rabbit (Saburkina et al., 2014), guinea pig (Hardwick et al., 2014; Steele et al., 1994). Some of these prior efforts provided limited mapping. Others represented gross staining of neurons with coarse-grain graphical representations of the areas involved. While qualitative and gross anatomical descriptions of the ICN have been presented earlier, we present here the first comprehensive neurocardiac atlas of the ICN in rat at cellular and molecular levels at a microscopic level.

The present datasets are a revelation of previously unsuspected complexity and diversity of modulators, receptors, and neurotransmitters that is already stimulating new anatomical and functional studies. The present discovery of phenotypical spatial gradients will stimulate connectomic studies to associate these with cardiac targets and functions going forward. Such work will ultimately be important for cardiac electrophysiologists performing ablation procedures. Prior work could not and did not relate cell scale neuroanatomy and cardiac scale organ anatomy, which our present results do. There is no prior literature on ICN transcriptomics or localization. This will be invaluable to autonomic nervous system investigators, vagus nerve cardiac regulation investigators, vagus nerve therapy investigators, cardiologists, heart anatomists etc.

## RESULTS

### Multi-disciplinary Approach to Data-acquisition Pipelines

We developed and applied a dual method pipeline to create a comprehensive anatomical map and molecular profile, within the 3D structure of the heart, of cardiac neurons in a rat heart. These integrative datasets are informative and useful in their own right as a first appreciation of the rodent ICN distribution and of single-neuron transcriptional profiling in ICN neurons. We combined a diverse set of technologies that enabled a geographically distributed network of researchers with distinct skills to coordinate and enable the present results. These data acquisition approaches are graphically represented in the two pipelines in Figure 1. In the pipeline represented in Figure 1A we used Knife-Edge Scanning Microscopy (KESM) in order to image and retain the 3D structure of the heart. In parallel, custom software was used to assemble and compress the resulting ~750,000 individual images in order to generate a single image volume. Annotation of the resulting image for mapping the ICN in histological context of the whole heart was performed with novel software created for this purpose (Tissue Mapper, MBF Bioscience, Williston, VT)

**Figure 1.**
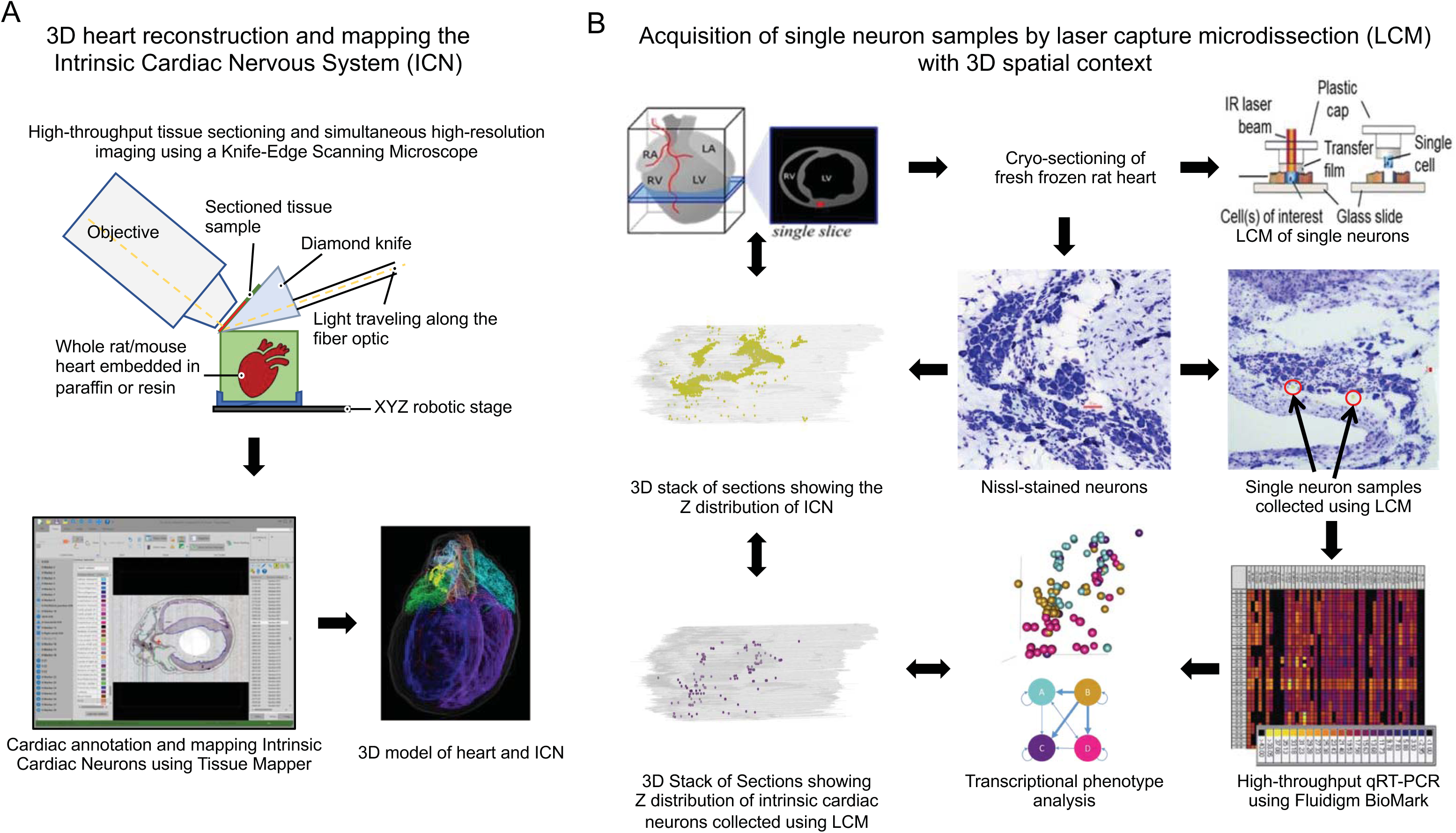
Data Acquisition Pipelines. (**A)** Acquisition of a 3D, accurate organ reference framework using high resolution collection of histological tissue sections. Once the entire heart is sectioned and imaged, the images are then compiled into an image volume by the TissueMaker software to enable 3D heart reconstruction and ICN mapping with the TissueMapper software. (**B)** Acquisition of cresyl violet stained neuronal samples from fresh heart tissue by cryostat sectioning. Single neurons were identified by position in the ICN and lifted for qPCR or RNA-Seq molecular phenotyping.

A second pipeline demonstrates the acquisition of neurons as single cell scale samples after cryostat sectioning of the heart to ascertain circuit connectivity and molecular phenotypes using transcriptional profiling (Figure 1B). Then the molecular phenotype(s) of these neurons can be placed in whole heart and ICN anatomical context. Images of these sections, including neuronal positions, are registered and aligned using Tissue Maker (MBF Bioscience, Williston VT) to create a whole heart volume with neuronal phenotype data that can be brought into the 3D reference system created by the first approach thus generating single-cell transcriptomics in a robust anatomical context that can incorporate data from multiple subjects for future comparison.

### Comprehensive Neuroanatomy of the Rat Heart

Heart tissue sections were imaged as they were cut using a 4th generation KESM platform (Strateos, San Francisco, CA), preserving tissue section morphology and registration to precisely conform to that of the intact, paraffin embedded organ. The image resolution (0.5 µm per pixel) yields clear visualization of cellular/neuronal scale histology. The registration and stacking of all the section images in a format compatible with the purpose-built software, TissueMaker and Tissue Mapper, supports using the latter to precisely map the position of each neuron and histological features of the heart and blood vessels from each tissue section. We then combine the positions of all neurons and features of cardiac tissues using computer graphics to hold all the data as a precise 3D representation of that specific heart’s ICN distribution in the histological context of the heart. The representation is precise and reproducible since both the tissue image acquisition and the mapping of neurons and cardiac features are under sub-micron control. This data acquisition pipeline is presented in Figure 1A. The complete data set of anatomical mapping is available online via SPARC Data Portal (http://data.sparc.science).

Since we take such care to represent the exact positions of all the neurons in a precisely preserved heart-organ morphology, we now can interact with the 3D model presenting all the mappings in various formats, scales or representations to appreciate the neuroanatomy of the ICN, enabling the following analyses.

- As illustrated in Figure 2 the ICN distribution is seen in the context of the whole 3D heart, depicting the unexpectedly extensive distribution of neurons on both posterior atria extending in the superior-inferior dimension from the base of the heart to the coronary sulcus or atrioventricular groove. **Figure 2.**
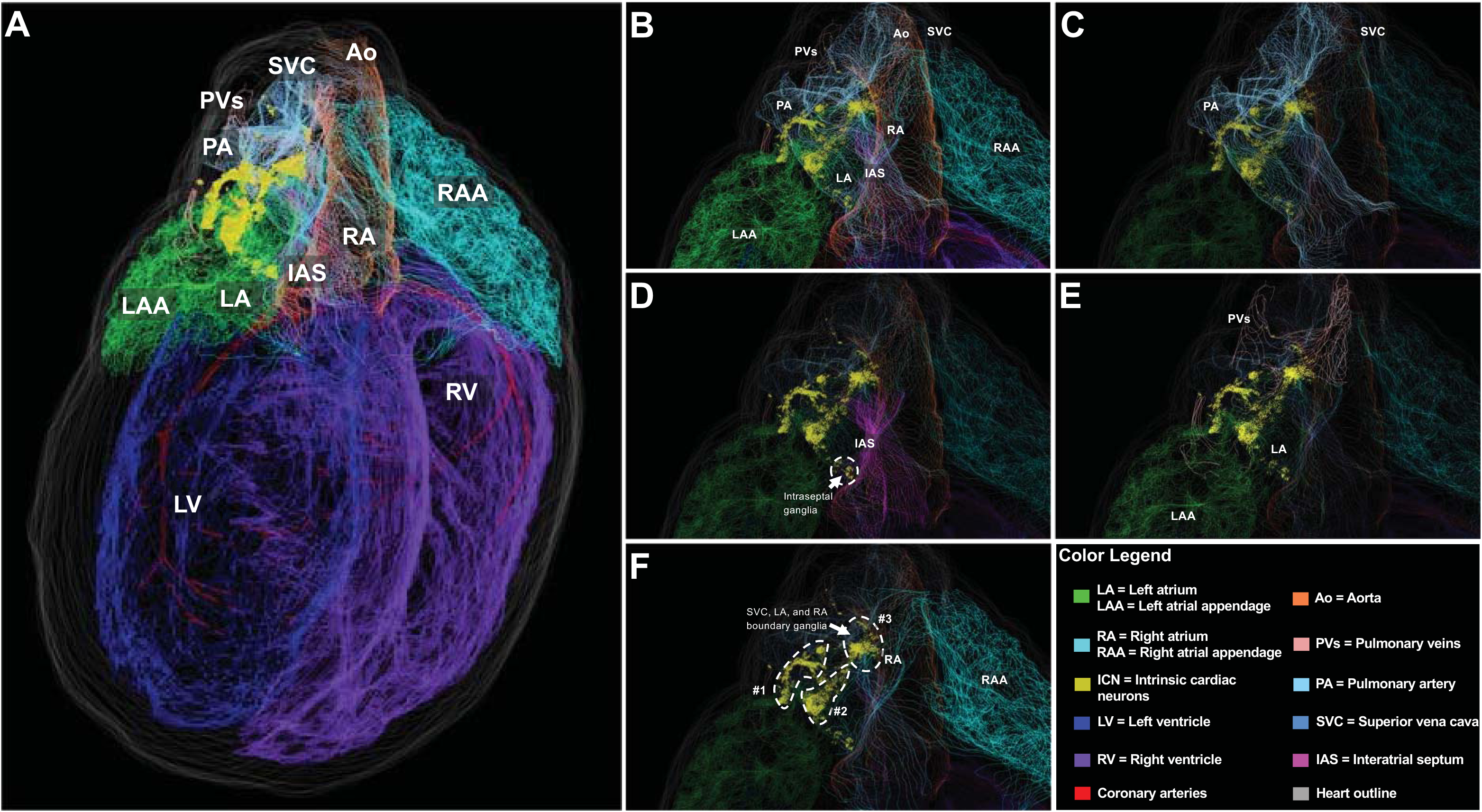
Posterior view of the 3D reconstructed male rat heart. **(A)** Whole-heart view showing the context, extent and distribution of the intrinsic cardiac neurons (ICN), located on superior and posterior surfaces of the atria. (**B-F)** take a higher-resolution view of the atria and blood vessels which are shown in **(B)**, with various contoured features of heart anatomy that are selectively removed to appreciate the anatomical relationship of the ICN to **(C)** the pulmonary artery and superior vena cava (SVC), **(D)** the interatrial septum, **(E)** the left atrium and pulmonary veins, **(F)** the right atrium, where clusters #1 and #2 appear to be on the surface of both atria, while cluster #3, located around the border of the superior vena cava, left atrium, and right atrium appears on the right atrium.
- We also are able to view subsets of the ICN in their local heart context by restricting the 3D model using the “partial projection” software tool. For example, Figure 3 is a 3 mm sagittal section representing neurons along the superior-inferior extent of the heart. Figure 4B creates a 3 mm thick section in the transverse plane that visualizes neurons located on the base/hilum of the heart. Whereas Figure 5 excludes these neurons showing only the more inferiorly positioned neurons. **Figure 3.**
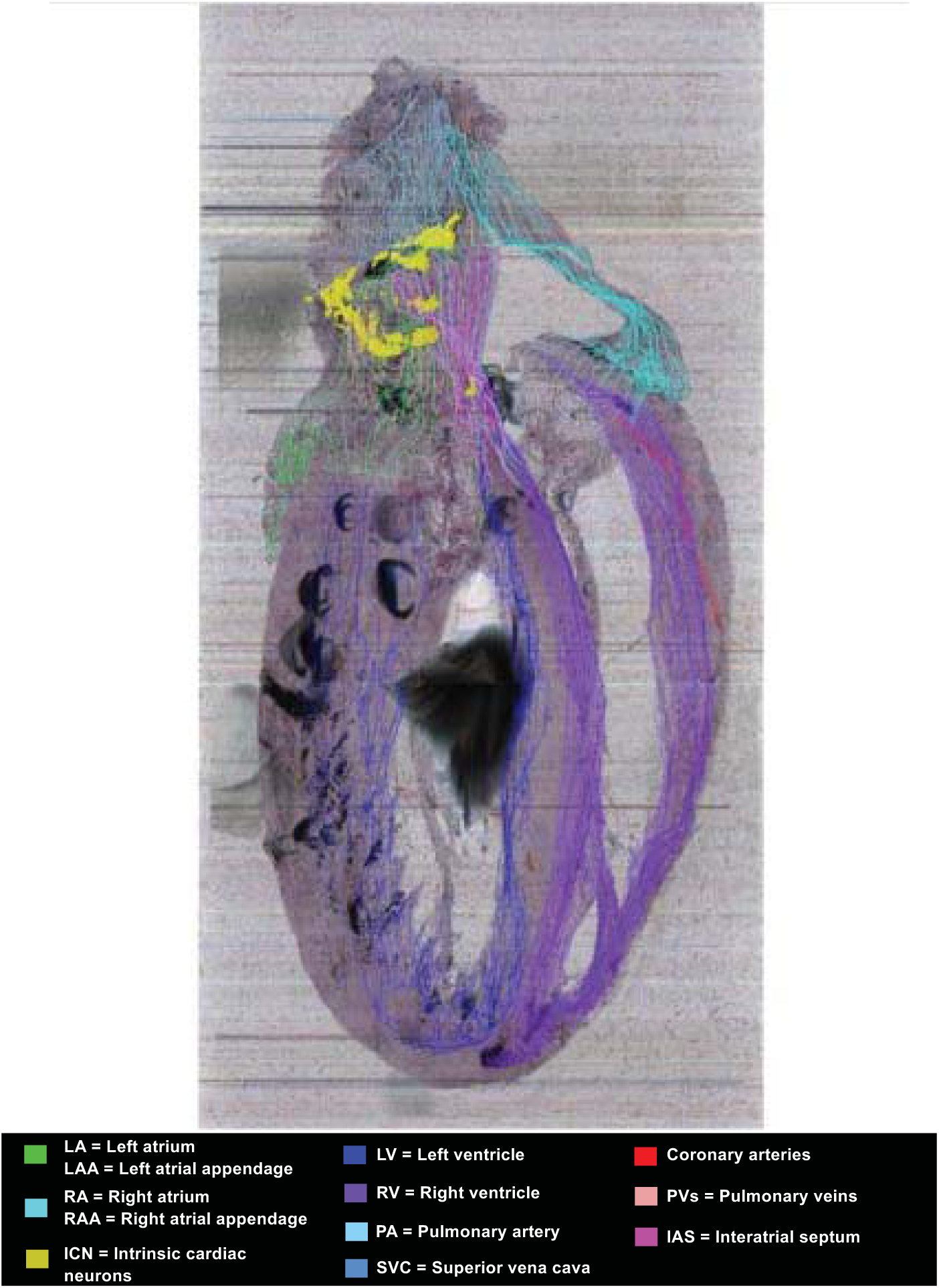
ICN distribution in a partial projection of sagittal heart sections. A partial projection of contours and ICN in a 3mm thick sagittal image slab illustrate the distribution of neurons along the superior-inferior extent of the heart.

**Figure 4.**
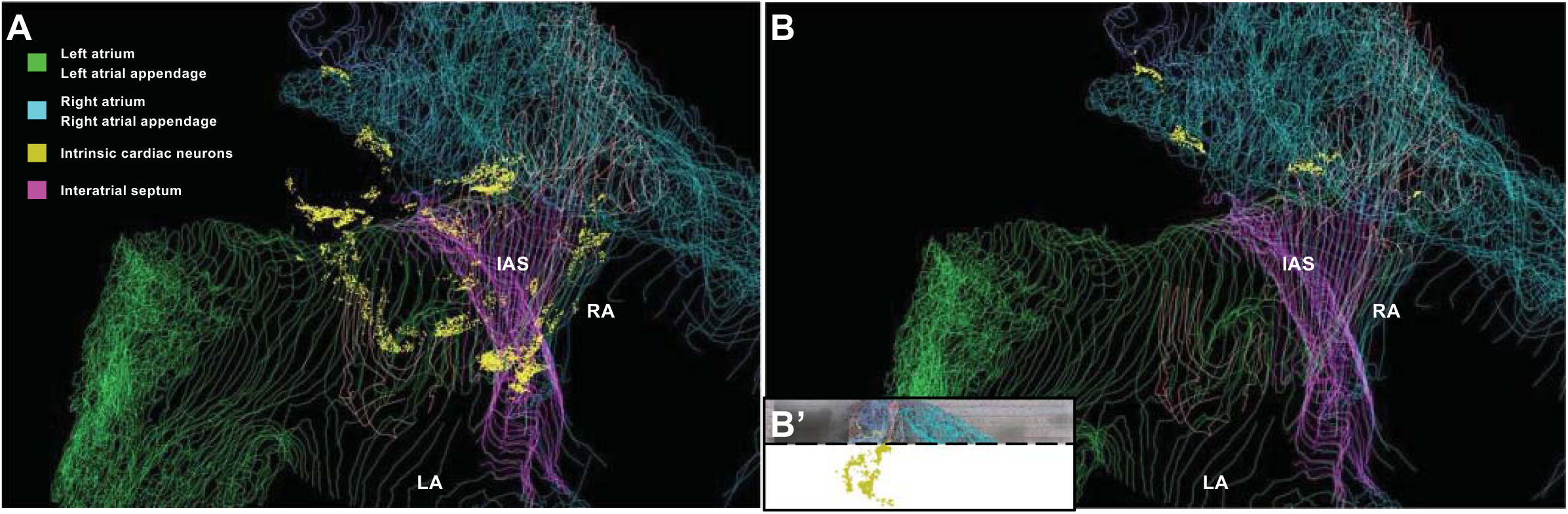
Superior view of the ICN. **(A)** Viewing the ICN distribution (neurons mapped as yellow dots) looking from the base towards the apex. The left and right distribution of all ICN neurons is discriminated by their relationship to the interatrial septum. Note that most ICN neurons visualized in **(A)** are not all on the base of the heart but mostly distributed at more inferior-caudal levels of the heart on both atria. **(B’)** In order to selectively view those neurons on the base of the heart we took this posterior view of the full ICN and retained only those above the cut-off point indicated by the dotted black line. **(B)** Then these neurons are here observed from the superior view of the heart, showing the position of ICN neurons located within the hilum in between the aorta, superior vena cava, pulmonary artery.

**Figure 5.**
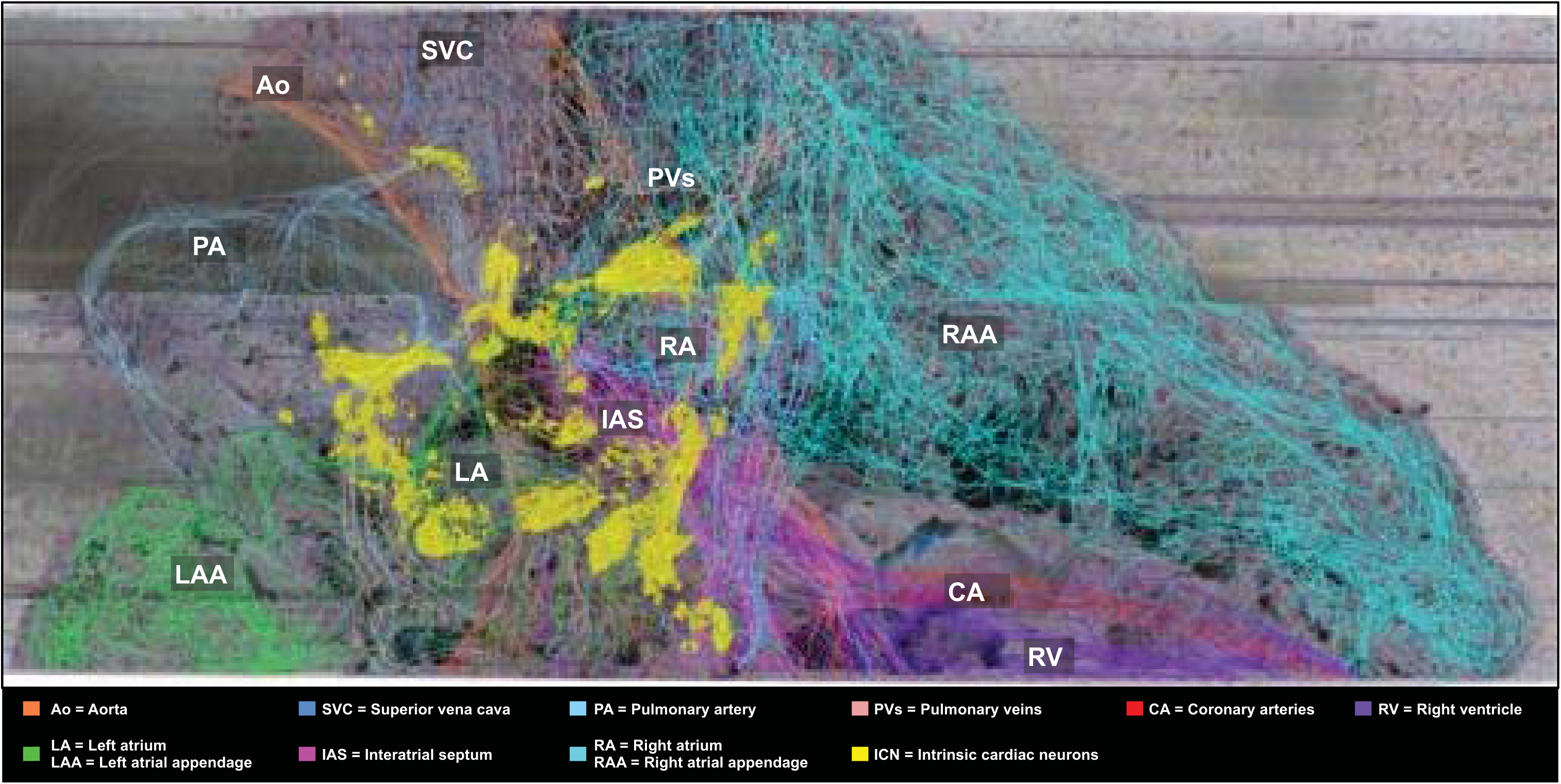
Distribution of more inferiorly located ICN. Using the TissueMapper Partial Projection tool that visualizes the ICN in a 3mm thick transverse image slab rotated in a superior view. This illustrates the locations in the transverse plane of section to highlight more inferiorly (caudally) located neurons. Neurons are represented as yellow dots.
- We are able to describe the exact position of ICN neurons in histological sections. Figure 6 consists of single sagittal section views of neurons throughout the superior-inferior extent in a series of sections going from right to left. **Figure 6.**
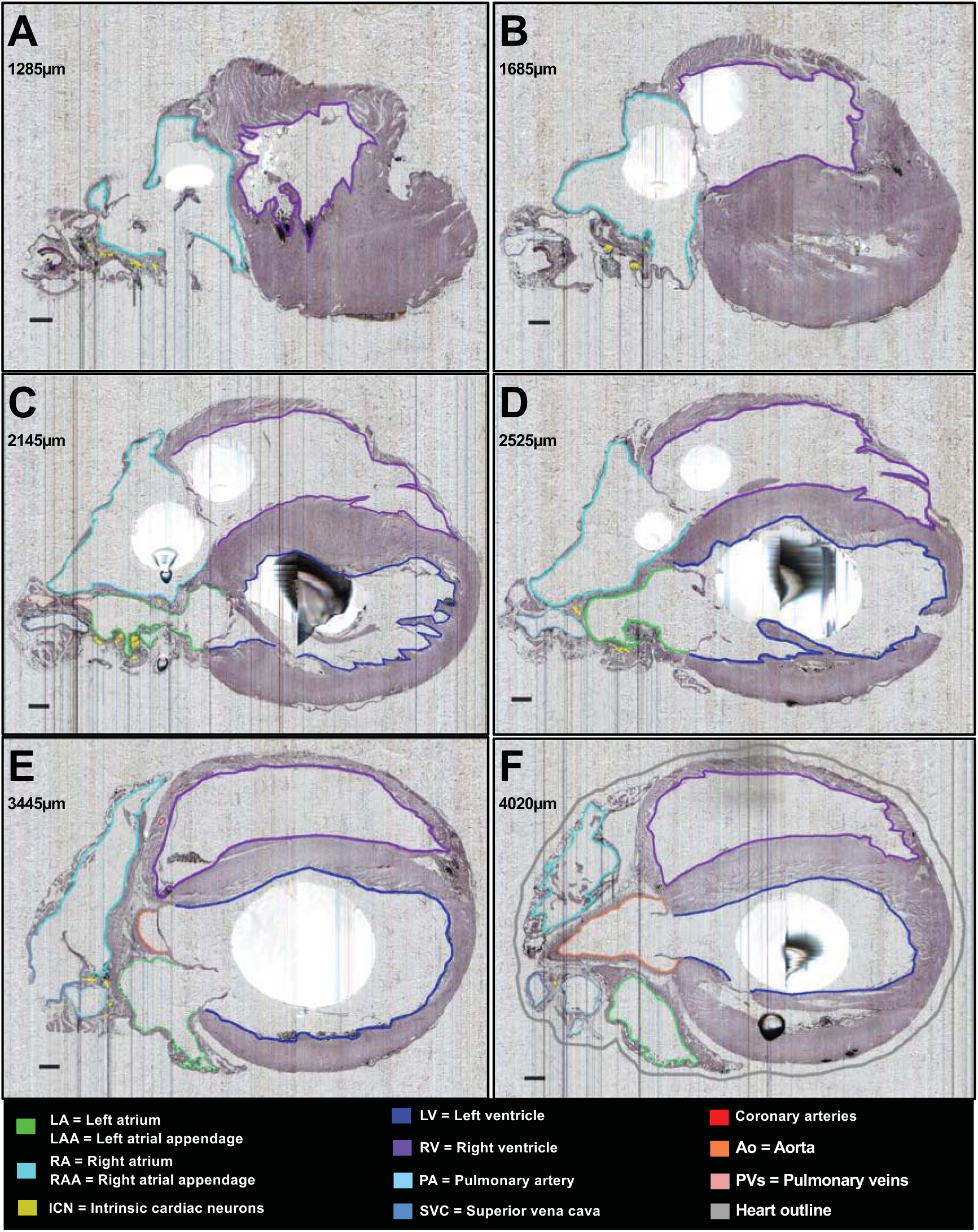
ICN distribution at six different semi-sagittal levels of the heart. **(a-f)** Histological sampling of the sagittal sections extending from right heart to left heart at the levels indicated in microns in each panel. The neurons are mapped with yellow dots. The contours help to contextualize the distribution of ICN relative to other features of the heart. Sections are 5um, images are at 0.5um X-Y resolution. The black blobs are artifacts of uneven paraffin embedding. Scale bar: 500µm.
- The digital neuron mapping supports quantitative analyses of neuron distributions and packing density as illustrated in Figure 7. **Figure 7.**
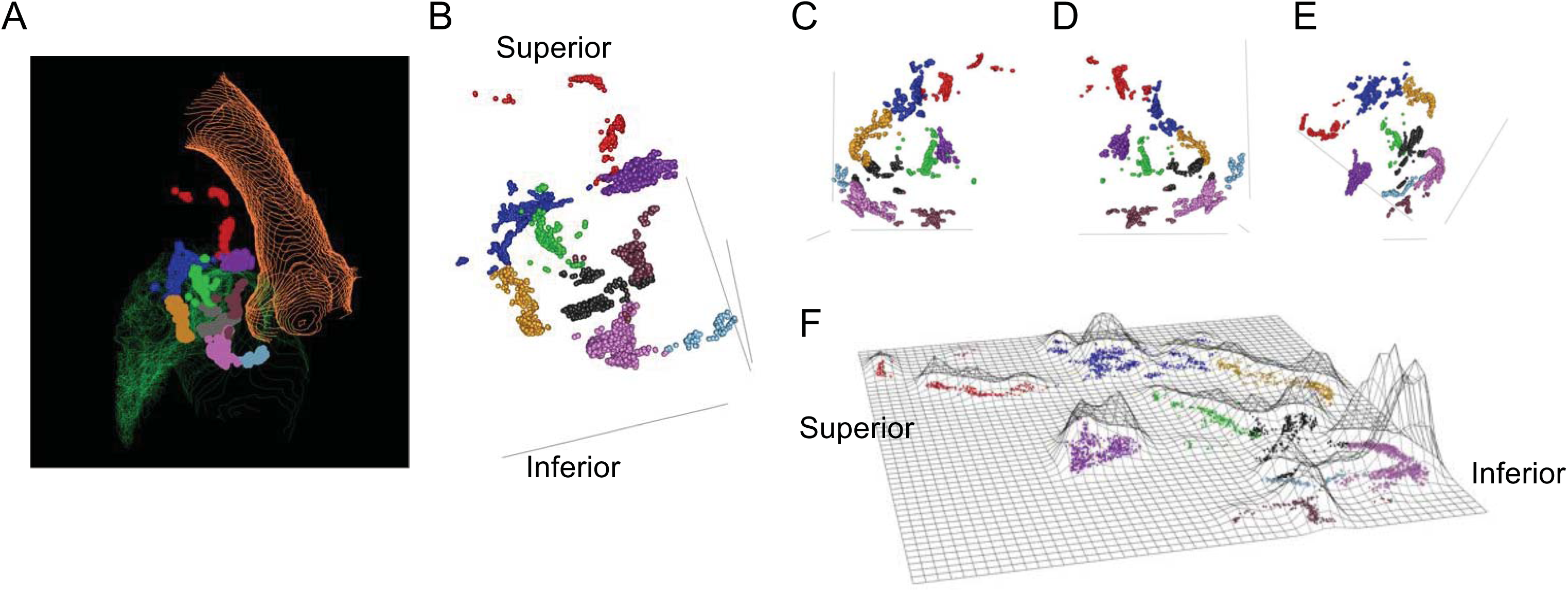
Identification of Neuronal Clusters in the Rat heart. Nine clusters of neurons were identified using the partitioning around medoids (PAM) algorithm, where identified clusters are shown in different colors. **(A)** Visualization of mapped neurons in their 3D orientation in TissueMapper, contours show the Aorta (orange) as well as the Left Atrium (green). **(B)** Visualization of mapped neurons in their 3D orientation. **(C-E)** Visualization of mapped neurons in their 3D orientation rotated to show different points of view. **(F)** Flat-mount projection of mapped neurons where the height of the contours are proportional the density of neurons. The orientation in (E) matches the flat-mount projection in (F).
- These anatomical templates above support addition of connectome and molecular identity data. Figure 8 shows the neurons in the digitized high-resolution images of histological sections. Figures 8 & 9 illustrate the strategy used in which we employed the pipeline represented in Figure 1B, to laser-capture single neurons for molecular analysis. Figure 10 visualizes the mapped ICN distribution to the distribution of neurons sampled and analyzed for molecular phenotypes. Figures 11-13 present analyses of molecular neuronal phenotypes, and demonstrate the gradients and distribution of phenotypes across the extent of the ICN. **Figure 8.**
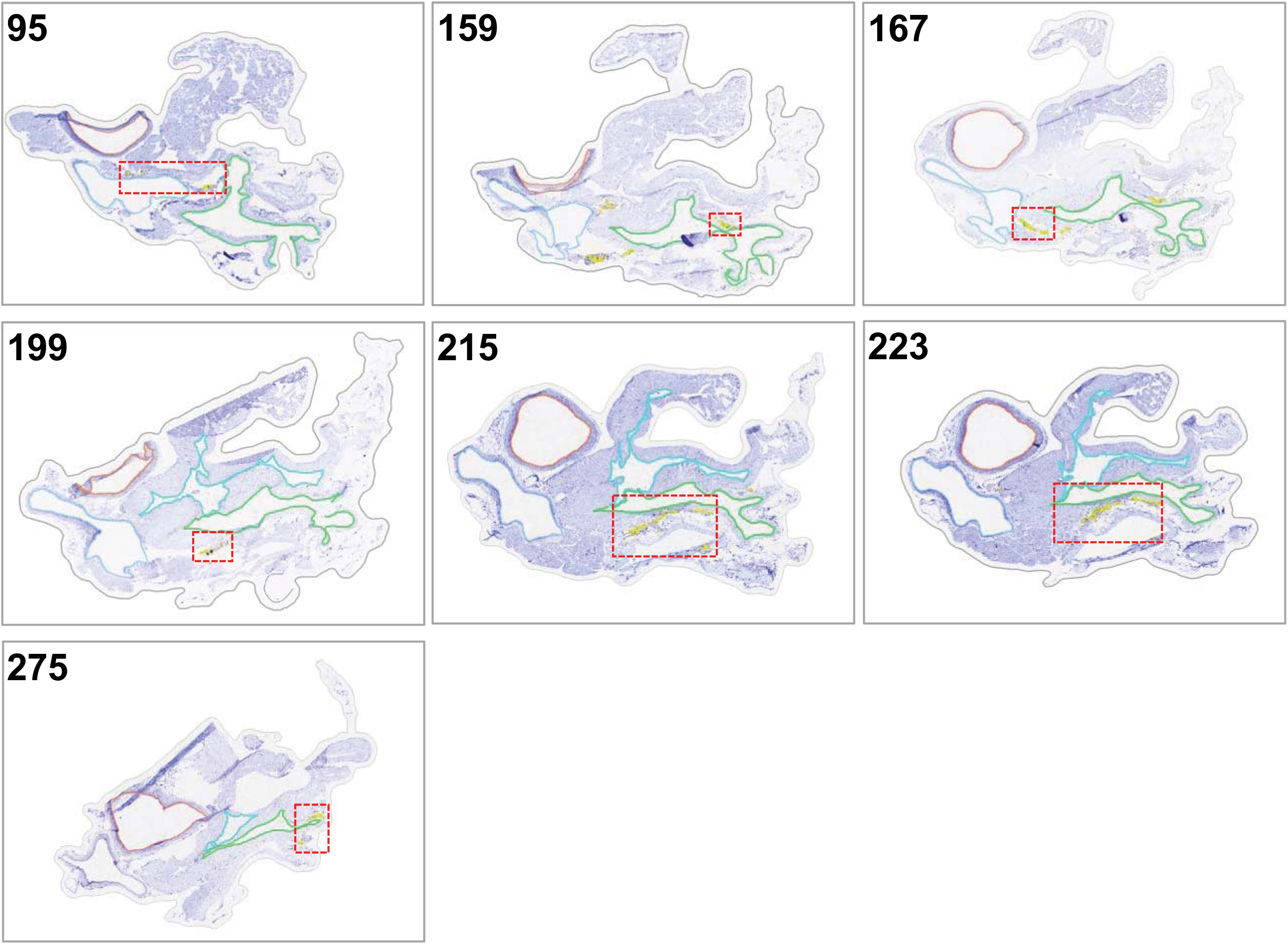
Candidate sections of female rat heart B to contextualize the laser capture microdissected intrinsic cardiac neurons. Each of the candidate sections provide additional context for the 7 levels of laser capture microdissected neurons. The red boxed areas are the regions of interest were zoomed in the following figure.

**Figure 9.**
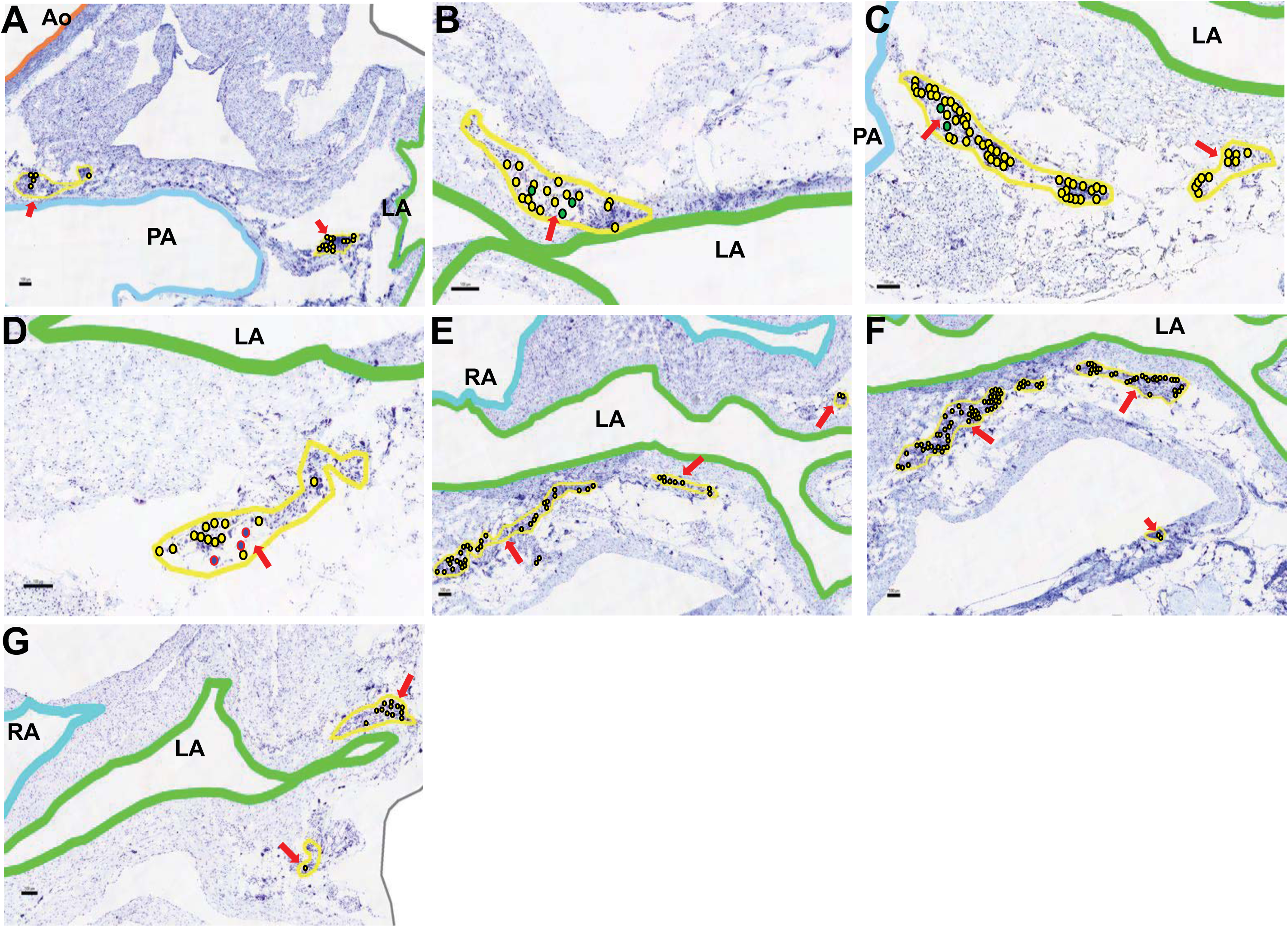
Enlarged sections of LCM lifted intrinsic cardiac neurons from 6 different levels in the female rat heart B. **(A)** Section 95-Small ganglia located between the aorta, pulmonary artery, and left atrium – all neurons marked yellow. **(B)** Section 159 – neurons are lifted from a medium ganglia near the left atrium and are grouped in the Z1 molecular cluster – cells marked green. **(C)** Section 167 – Z1 neurons that are located around the left atrium and pulmonary artery – cells marked green. **(D)** Section 199 – LCM sampled neurons that were characterized as the Z2 group were part of a small ganglia – cells marked blue. **(E)** Section 215 – neurons were sampled from ganglia near the left atrium and right atrium. **(F)** Section 223 – LCM sampled neurons from a large cluster near the left atrium **(G)** Section 275 – LCM sampled neurons around the left atrium. Scalebars: 100µm. Red arrows mark the locations of relevant neurons.

**Figure 10.**
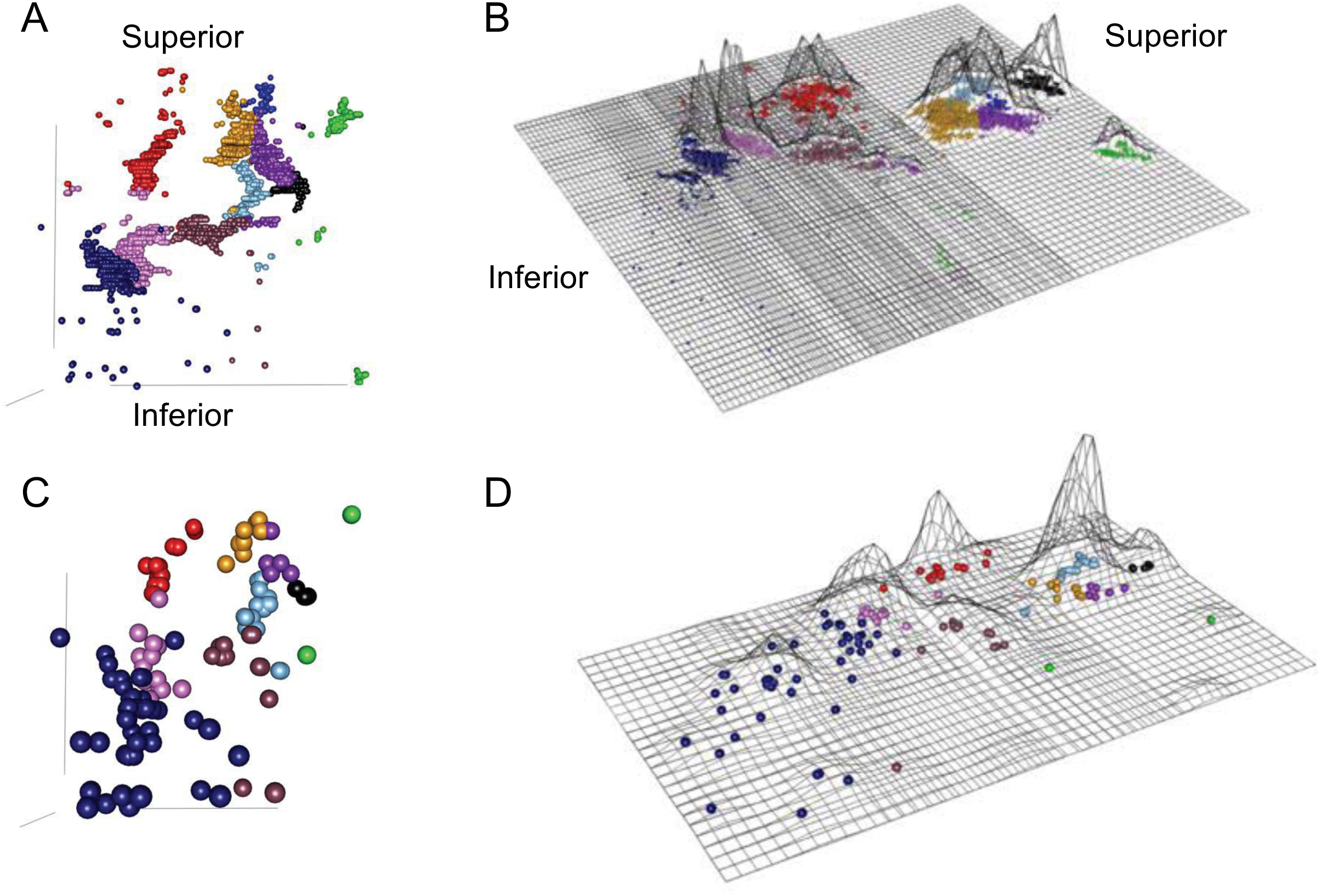
Identification of neuronal clusters in the Rat Heart—Heart B. Visualization of mapped neurons **(A)** and sampled neurons **(C)** in their 3D orientation. Flat-mount projection of mapped neurons **(B)** and sampled neurons **(D)** where the height of the contours are proportional to the density of neurons. 10 clusters of neurons were identified using the partitioning around medoids (PAM) algorithm, where identified clusters are shown in different colors.

**Figure 11.**
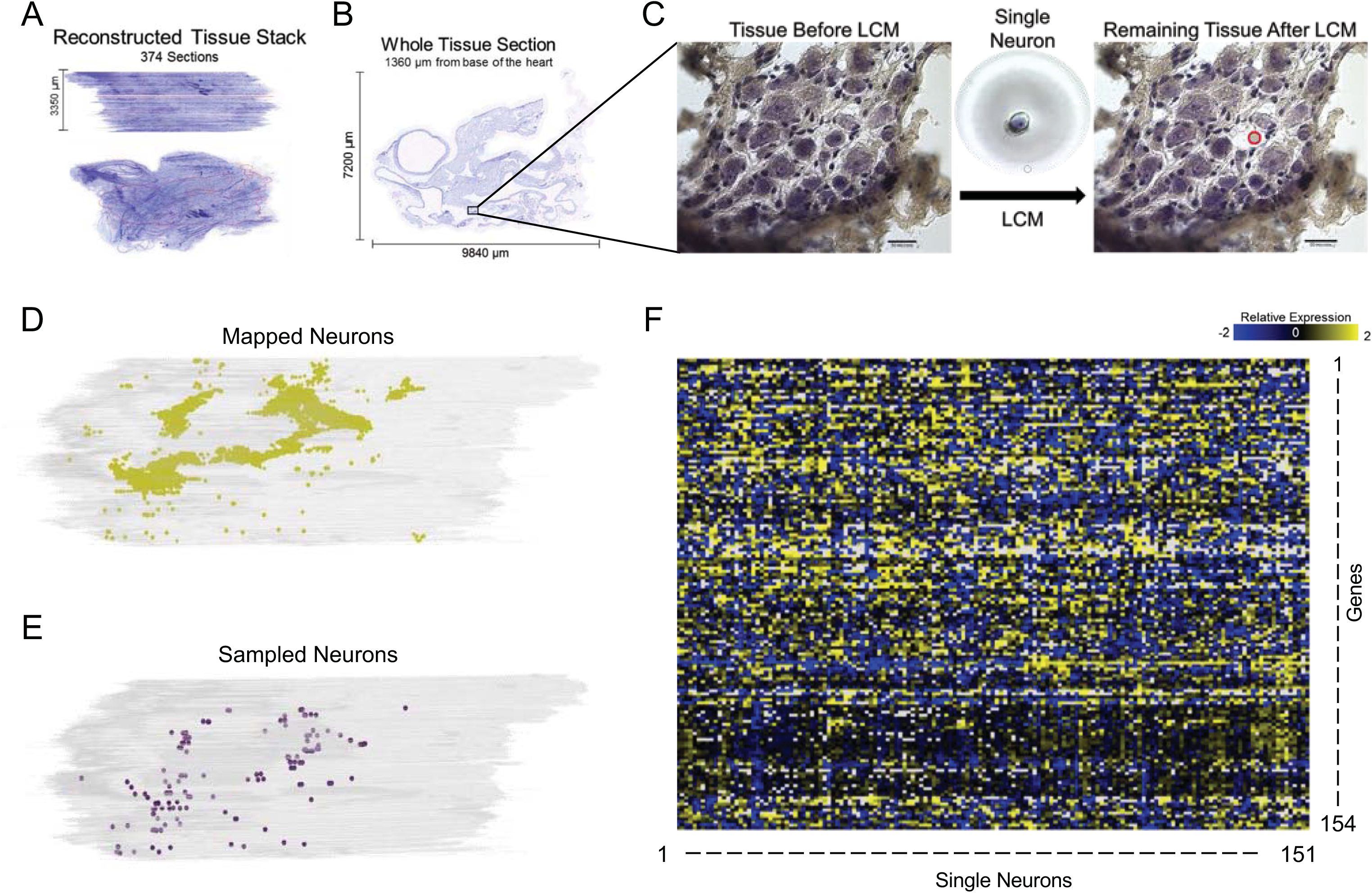
Process of collecting and mapping Neurons from the rat heart. **(A)** A rat heart was sectioned at 20 um, imaged, and put into a 3D stack in TissueMaker. The red outline shows a specific section shown in (B). **(C)** the selected region from **(B)** is shown in a magnified view before and after laser capture microdissection, where the single neuron that has been collected can be seen on the LCM cap (middle). Distribution of mapped neurons **(D)** and sampled neurons **(E)** in the context of their 3D location as seen in TissueMapper. **(F)** Normalized qRT-PCR data showing expression of 154 genes for the 151 samples collected.

The results from these visual and quantitative analyses are presented below in detail. In addition to the figures, the visualizations are also represented in movies available in the Supplemental material.

We sought to delineate the distribution, extent and location of neurons on the heart in part by the use of 3D graphical models of the ICN in cardiac context both in figures and by rotating the model (Supplemental Video 1). Examining the 3D model framework of the ICN from a posterior point of view, shows the entire population as compact and localized to a region on the superior and posterior aspects of both left and right atria, extending inferiorly to the atrioventricular boundary or the coronary sulcus at the boundary with the ventricles, as seen in Figure 2A. The additional panels, Figure 2B-F, are enlarged images of select regions of the heart to highlight the anatomical features of ICN, in each of which various contours outlining specific features of cardiac anatomy are dimmed. This provides a better appreciation of the ICN distribution, shape, extent and location in relation to specific cardiac structures. With all contours illuminated the zoomed-in ICN appears to comprise three or more large clusters in which neurons extend rostral to caudal almost continuously (Figure 2B). At its most superior extent portions of the ICN lie adjacent, semi-surrounding and posterior to the pulmonary artery and superior vena cava, on the right side of the heart (Figure 2C). As neurons extend Inferior to these levels they distribute in the shape of “a comma” and extend laterally to the left. Neurons at these levels are situated on both atria and within the interatrial septum (IAS) (Figure 2D). Further inferior the ICN neurons can be seen to distribute in relation to the contours of the pulmonary veins in Figure 2E, extending on the posterior surface of the left atrium to the boundary with the left ventricle. In summary, the 3D reconstruction illustrates how the most prominent clusters of neurons are positioned on the left atrium and/or across both atria as in Figure 2F. Most neurons appear to be located on the posterior surface of the left atrium. Other neurons, more superiorly located, are located within the interatrial septum and on the right atrium. This again shows that the ICN is associated with the posterior wall of both left and right atria as well as the interatrial septum (Cheng et al. 1999, Ai et al. 2007b, Cheng et al. 2017). Using visual inspection for grouping, there are three large clusters of ICN neurons associated with the atria denoted as #1, #2, and #3. From this perspective, Clusters #1 and #2 appear to be principally on the epicardial surface of the left atrium, while cluster #3, located adjacent to the borders of the superior vena cava, left atrium, and right atrium, appears to be largely on the right atrium.

The ICN distributes on the posterior atria. In order to position, in higher resolution, the neurons associated with posterior cardiac structures, we used the virtual section “partial projection” tool from the Tissue Mapper software. Using partial projection, a user-defined subset of the 3D heart model and neurons in any plane and thickness can be displayed, more than single sections and less than the full 3D model. Using this tool we made a virtual section that displays only neurons in a 3 mm sagittal section oriented superior-inferior, centered on the posterior cardiac surface, as seen in Figure 3. This shows a more restricted portion of the heart structure while presenting a more coherent view than single sections can provide. At these levels the majority of neurons appear inferior to the base of the heart, extending to the coronary sulcus. This thick section and point of view is focused on the posterior surface of the heart. This has the advantage of providing a broader context than single sections but also a more focused examination than the full 3D heart context. In Figure 3 this partial projection of the ICN localizes a compact population of neuronal clusters with a superior-inferior orientation, distributed on both atria and in the IAS, but preferentially on the left atrium.

By rotating the 3D model, we found that a superior view, i.e. looking down at the base of the heart, provided the ability to distinguish neurons associated with the right vs. the left atrium. Figure 4A takes a superior view of the 3D heart and ICN, viewing the base of the heart from a viewpoint that highlights the profile of the IAS. In particular, the distribution of neurons in the left and right atria can be observed in relation to the IAS, with neurons associated with both atria, but more so to the left atrium, with a much smaller population within the IAS itself. In Figure 4A, the entire population of the ICN is seen from a superior viewpoint allowing appreciation of the full distribution of neurons on the left and right atria.

Since in rat the ICN is often associated with the hilum on the base of the heart we examined only those neurons with that specific neuroanatomical distribution and localization. In Figure 4B’ and 4B we aim to acquire and then only visualize neurons plausibly located on the base of the heart. As described above, using “partial projection”, a user-defined subset of the 3D heart model and neurons in any plane and thickness can be displayed, more than single sections and less than the full 3D model. Using the partial projection tool, we made a virtual section that displays only neurons at the base of the heart (as seen in Figure 4B). These ICN neurons are on the epicardial aspect of the base associated with the major pulmonary vessels there, the cardiac hilum. These neurons appear to be on the right atrium and in the interatrial septum.

Our results show that the majority of neurons are positioned more inferiorly to the base of the heart extending caudally to the boundary with the ventricles as seen in Figures 2 & 3. In order to visualize their distribution and location across the two atria we again took a superior view of the heart to observe left-right neuronal distribution of neurons, but used the Tissue Mapper software “partial projection” tool to “cut off the base/hilum) and observe only neurons inferior to the base of the heart (Figure 5). This graphic shows those ICN neurons as a transverse or axial visualization of a 3 mm thick slab of tissue. Neuron clusters are abundant across both atria, but in contrast to the base of the heart neurons are more abundant on the left atrium.

The 3D reference models of the heart and ICN are derived from whole-mount stained histological tissues (i.e., whole rodent hearts). It is also informative to look at these single tissue sections and appreciate the position of the neurons in them. The “native” section images in Figure 6 are from the data acquisition pipeline in Figure 1A. These six tissue section images are at 0.5 µm (X-Y) resolution and allow appreciation of the cellular histology for discriminating neurons. As shown at six different levels of the heart (Figure 6) individual neurons could be identified and their locations determined in relation to the heart chambers and other structures. This heart was sectioned in a near-sagittal plane and Figure 6A-F moves sequentially from the right to the left side of the heart. The image volume is available online through the SPARC Data Portal. Figure 6A is close to the right side limit to the ICN and 6F is close to the left limit. The numbers in each panel indicate the right-left position of the section. Thus, Figure 6A is 1285 micrometers from the right heart edge and Figure 6F is 4020 micrometers farther left. In these tissue sections, neurons form clusters on the surface of the walls of the right and left atria. As seen in Figures 2 and 4 these clusters continue across many sections, so that when the maps are combined (stacked) across sections, these neurons form somewhat continuous structures in the 3D model. These structures may represent previously identified cardiac ganglionated plexuses (Cheng et al. 1999, 2004).

Whereas the qualitative inspection of ICN clustering finds areas of high and low neuronal packing density, a more quantitative and objective approach to these observations involves use of “partitioning around medoids (PAM)” algorithm to assay packing density or clustering (Kaufman and Rousseeuw, 1987). These clusters can be viewed in Tissue Mapper in 3D cardiac context (Figure 7A; Supplementary Video 2) and without cardiac features, as the ICN alone, in a posterior point of view (Figure 7B) and, in Figures 7C-E rotated points of view to highlight the shapes of the clusters. These clusters can be represented as surface plots, as in Figure 7F where the height of the surface contour represents the neuronal packing density.

It is possible that, like the neurons in the brain, these clusters of neurons may be somewhat variable in precise spatial distribution from animal to animal, but identifiable as they are consistently present at the coarse-grain anatomical locations relative to notable cardiac features. By analyzing the connectivity and molecular phenotype of individual neurons from these clusters, it may be possible to identify functional groupings within the ICN. They may have a consistent connectome of functionally specific projections, and may also have distinct molecular phenotypes. The representation of the anatomy of molecular phenotypes of the ICN requires the data acquisition pipeline described in Figure 1B. Figures 8 & 9 show rat heart sections acquired in this pipeline. Figure 8 presents overviews of regions containing ICN neurons from which LCM samples were acquired, and Figure 9 the detailed neuronal groupings from which single neurons were sampled.

### Combining 3D anatomical mapping of the ICN with laser capture microdissection of single neurons for molecular phenotype analysis

The sections in Figures 8 & 9 are single sections from a female rat heart in which we mapped the 3D organization of the ICN. The heart was sectioned in the transverse plane going rostro-caudally between the base and apex. The tissue sections shown are at regular intervals at levels where ICN neurons are present in the heart. At each level the position of captured neurons is indicated in yellow. These distributions in two dimensions are like what is seen in Figures 2–5 in 3D or stacks, with relatively continuous distributions of neurons that tend to clump in groups consistent with being described as ganglionated plexuses. Distinct clusters of widely separated cell groups are associated with both atria. Sections are numbered from the base and neurons were found in sections 39 through 470.

The neurons marked with yellow dots in Figure 9 indicate the single cells mapped within the ICN. An effort was made to select cells randomly but broadly representative across all sections. Neurons that were lifted from the tissue by performing laser capture microdissection (LCM) at Z1 levels are marked green (Figure 9B & 9C) and those from Z2 levels are marked in blue (Figure 9D). Such microdissected neurons (n=151) were used to perform single cell transcriptomics. The molecular phenotype of each neuron is associated with the position of each cell in the heart’s 3D coordinate framework. The genes expressed and the cell types of these isolated neurons combined back into the specific distributions in 3D context generated by stacking the serial images. This quantitative display can then be used for the visualization of laser capture neurons and linked to their molecular phenotype datasets. Figures 10A & 10B uses the “data driven” ICN quantitative clustering as was done in Figure 7. However, this ICN was “undersampled” for mapping and is not for comparison to Figure 7. Rather it is to provide context for the clustering and distribution of the ICN neurons lifted for molecular profiling in Figures 10C & 10D.

### Molecular analysis of single neurons in the 3D ICN context

We obtained a high-throughput transcriptomic data set containing 23,254 data points from 151 samples of single neurons (Figure 11). Each sample was assayed for the expression of 154 genes selected as associated with neuromodulation and cardiac function. We analyzed this transcriptomics data using Principal Component Analysis (PCA) to identify the structured variation that, if present, could organize the neurons into subgroups based on the variability in the gene expression profiles (Figure 12; Supplemental Video 3). We mapped the three major neuron groups that arose from PCA results to their spatial location in the three-dimensional coordinates of ICN. Interestingly, the superior-inferior position (base to apex direction) accounted for the most robust clustering along the first two principal components PC1 and PC2 that account for the top two dominant sources of variation in the gene expression data (Figure 12A). Samples were divided into three Z position groups (Z1, Z2, Z3) where Z1 is closest to the base, moving down to Z3 near the inferior aspect of the atria (Figure 12). Examining the gene expression profiles of samples within the Z groups, distinct sets of genes were distinctly enriched in each of the Z1, Z2, and Z3 positional groups, with a small subset being enriched in both Z2 and Z3 (Figure 12C). Notably, the neurons within each of the Z groups were isolated from multiple ganglia (Figure 11), suggesting that the heterogeneity of molecular phenotypes is more constrained than the spatial distribution of these neurons.

**Figure 12.**
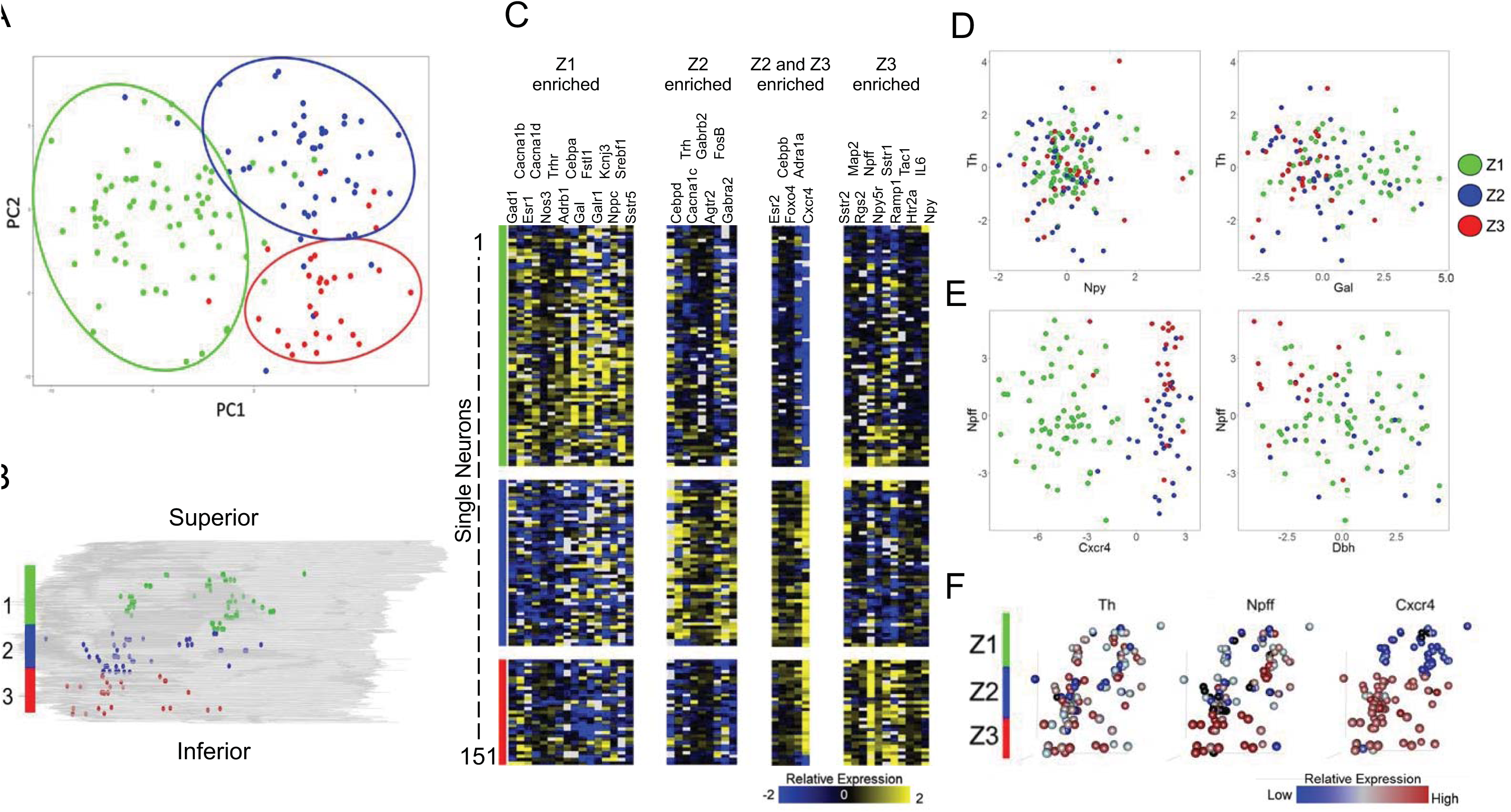
**(A – B)** Principal component analysis (PCA) plot of the 151 collected samples (A) show distinct separation according to their position along the Z axis of the heart (B). Samples were divided into three groups along the Z axis spanning from Z1 at the base of the heart to Z3 towards the apex. **(C)** Pavlidis template matching using Pearson correlation with a cutoff of 0.01 was used to find genes that show specific enrichment in one or more Z groups. A cutoff of 0.001 was used to find genes enriched in both the Z2 and Z3 group to increase specificity. **(D)** Expression of tyrosine hydroxylase (Th) vs. neuropeptide Y (Npy) (left) and Galanin (Gal) (right). **(E)** Expression of neuropeptide FF (Npff) vs. Cxcr4 (left) and Dbh (right). **(F)** 3D position of collected samples colored for expression of Th, Npff, and Cxcr4. While Th (F, left) and Npy (D, left) show little correlation between expression level and spatial location, Galanin (D, right) appears to be upregulated in the Z1 group. Npff (F, middle and E, left) shows downregulation in the Z2 group and distinct upregulation in the Z3 group. Cxcr4 (F, right and E, left) shows very high correlation between expression level and spatial location where expression is significantly downregulated in the Z1 group and significantly upregulated in the Z2 and Z3 groups. Dbh (E, right) shows mixed expression in the Z1 and Z2 groups with low expression in the Z3 group. It can also be seen that Npff and Dbh are anticorrelated in the Z3 group. For all panels, Z1 is represented in green, Z2 in blue, and Z3 in red.

We also analyzed the distribution of select molecular profiles in a pair-wise fashion to assess the alignment of conventionally described cell types in the ICN. For example, a pair-wise comparison of Th and Npy expression demonstrated that while several neurons could be classified as Th only or Npy only, a subset of neurons were Th and Npy positive. These Th+ Npy+ neurons were distributed throughout ICN without any apparent bias towards a narrow spatial location (Figure 12D, left). These results are consistent with results from immunohistochemistry where NPY positive cells and TH positive cells were observed as broadly distributed in the ICN (Richardson et al., 2003), but did not have a correlated expression as was suggested in other results (Crick et al., 1994). Examination of Galanin along with Th, however, shows a more organized pattern with higher expression of Galanin in the Z1 position groups, but with little correlation with Th generally (Figure 12D, right). Interestingly, we found that certain genes showed a very strong spatial localization bias, and examining them in a pairwise manner reveals further co-localization patterns. For example, Cxcr4 and Npff were highly correlated with spatial location. Cxcr4 expression was almost exclusively high in the Z2 and Z3 positional groups, whereas Npff expression was highest in Z3, lowest in Z2, with mid-range of expression in Z1 (Figure 12D, left). Cxcr4 and Npff were co-expressed in the neurons in the Z3 positional group. By contrast, the neurons in the Z3 positional group with high expression levels of Npff showed distinctively low expression levels of Dbh, with little correlation between the two genes in Z1 and Z2 neurons (Figure 12D, right). Cxcr4 is also known as the NPY3 receptor, suggesting that these neurons may be responsive to NPY+ sympathetic fibers projecting to the heart from the stellate ganglion. Npff has been shown to increase heart rate when acting on receptors in the rat heart, working synergistically with adrenergic signaling pathways (Allard et al., 1995). Using the mapping techniques discussed above, we can also visualize the expression patterns of these genes within the context of their three-dimensional localization, making it possible for us to delve deeper into the relationship between their expression patterns and locations within the rat ICN (Figure 12F).

We evaluated the gene expression data for differences between the three Z positional groups using a one-way ANOVA. Hierarchical clustering of samples using the genes with ANOVA p<0.001 elucidated four subtypes of neurons (A, B, C, D; Figure 13), with subtypes A and C being found primarily in the Z1 group, B in the Z2 group, and D in the Z3 group along with a number of samples from the Z1 group (Figure 13A&B). We analyzed the molecular patterns that underlie the spatially organized neuronal subtypes to identify correlated modules of gene expression and the combinations of modules that distinguish each neuronal subtype. For example, we see that comparing Npff to Cxcr4 expression largely delineates the different phenotypes. We saw that Npff and Cxcr4 greatly correlate with spatial position of the Z groupings, where the Z1 group shows a range of expression of Npff. When examining phenotypes, however, we see that phenotype A is largely comprised of cells that have low expression of both Npff and Cxcr4 while phenotype C largely accounts for the cells which show high expression in Npff and low expression of Cxcr4. We also examined the distributions of neuropeptide and corresponding receptor gene expression across the four identified phenotypes. For example, expression patterns between galanin and its receptor, Galr1, show that most cells in the phenotype A express high levels of galanin, with a large subset also expressing high levels of Galr1, indicating that cells in phenotype A largely drive galanin signaling both through autocrine signaling as well as acting as the source for paracrine signaling while cells from phenotypes A, B, and C act as the target for paracrine signaling. Most cells in phenotype D showed low expression for both Gal and Galr1 (Figure 13D & 13E). Examining expression patterns between thyrotropin releasing hormone (Trh) and its receptor Trhr show very different cell-cell signaling connectivity patterns when compared with those for galanin. While a small set of cells from all four phenotypic groups show coexpression of both Trh and Trhr, the most noticeable pattern is that phenotype B has consistently high levels of Trh, but not Trhr, indicating that cells in phenotype B primarily act as the source for paracrine signaling between phenotype B and the other three identified phenotypes.

**Figure 13.**
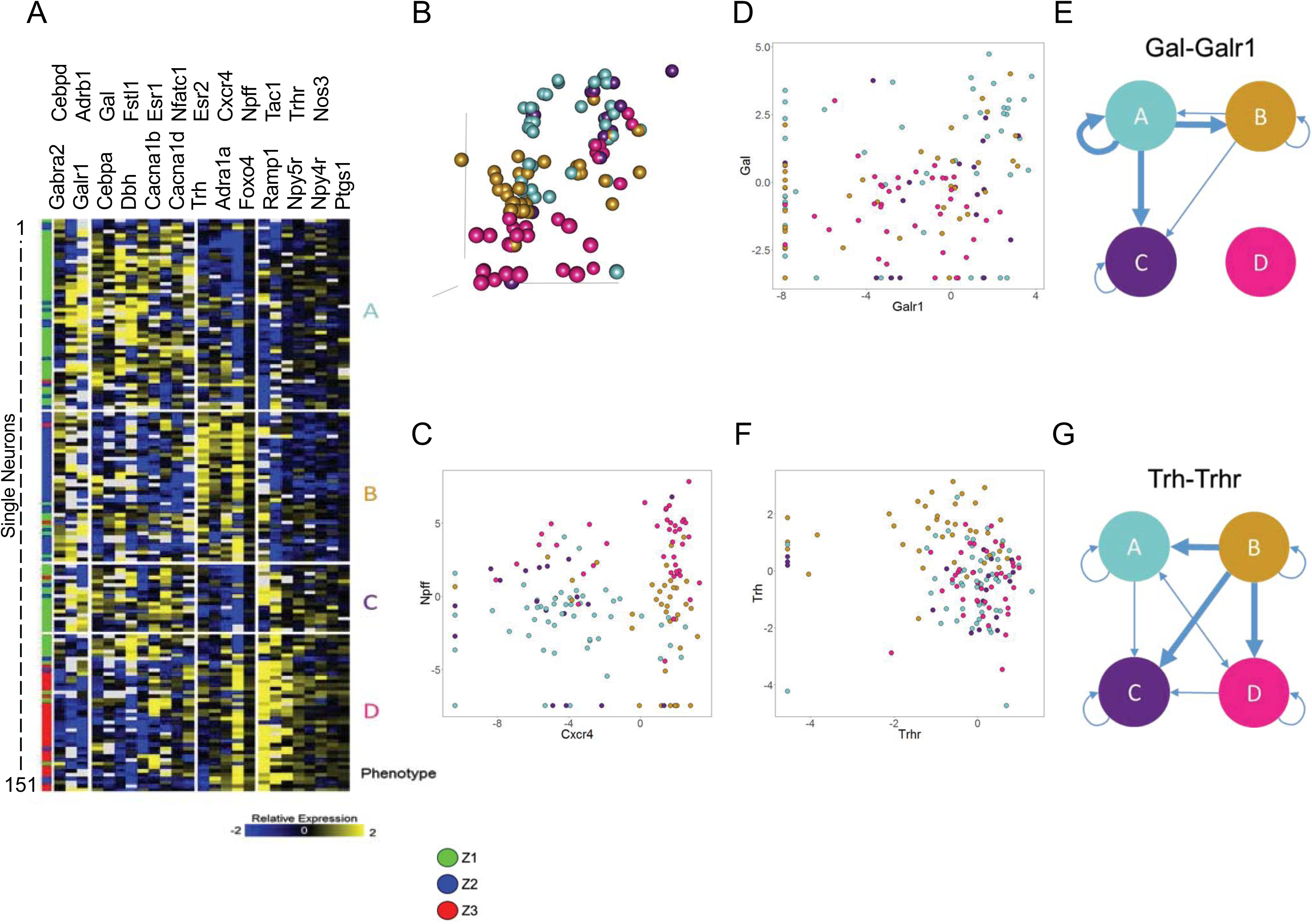
Neuronal Phenotypes contributing to Spatial Separation. **(A)** ANOVA was performed with a threshold of 0.001 to find genes that contribute to the separation seen between the three Z groups. Selected genes were clustered using pearson correlation, resulting in the heatmap shown and giving rise to four different neuronal phenotypes (Labeled A-D). We can see that the Z1 group was split into phenotypes A and C, with some samples from the Z1 group clustering along with the Z3 group in phenotype D. Phenotype B is comprised mostly of samples residing in the Z2 group. **(B)** 3D position of collected samples, colored by phenotype. **(C)** Expression of Npff vs. Cxcr4, colored for phenotype. We can see that the comparison of these two genes largely determines the phenotypes. Phenotype A comprises mostly of cells that have low expression of Npff and Cxcr4. Phenotype B comprises mostly of cells that have low expression of Npff and high expression of Cxcr4. Phenotype C comprises largely of cells that have high expression of Npff and low expression of Cxcr4. Phenotype D comprises largely of cells that have high expression of both Npff and Cxcr4. **(D)** Expression of Galanin vs. its receptor Galr1 and **(E)** network diagram of putative galanin-mediated connectivity between neuronal phenotypes. **(F)** Expression of thyrotropin releasing hormone (Trh) and its receptor Trhr and **(G)** network diagram of putative Trh-mediated connectivity between neuronal phenotypes.

## DISCUSSION

### Recapitulation of the major findings and their significance

In this paper, we present a method to identify individual neurons in the cardiac ICN and comprehensively record the neurons’ relative spatial positions in the heart at micron scale resolution for the first time. Combined with the ability to extract these neurons for downstream applications, we can now understand the gene expression of individual neurons in the ICN in their spatial context. Together, we present an integrated pipeline for cardiac connectome and molecular phenotype data acquisition. Here, we also demonstrate the first use of KESM (McCormick, 2002), on cardiac tissue for the purposes of making a cardiac atlas at a micron-level resolution. This process and approach can be expanded to other organ systems for the microscopic mapping of neuronal structures.

All these results were made possible by a team science approach involving four groups each providing distinct but necessary technology, as required by the SPARC Program. The approach demonstrated here should be applicable to other organs with intrinsic nervous systems. This approach can also be extended to human hearts in health and in heart failure, seeking neuromodulators involved in regulating heart function. Understanding the cellular and molecular processes governing innervation and the functional control of the myocardium in health and disease is the mechanistic basis for the development of neuraxial therapies to prevent sudden cardiac death and arrhythmias (Fukuda et al. 2015). The anatomical and molecular data are all archived in the DAT-CORE Portal of the SPARC Program for public access and use. The inspiration for our approach comes from the Allen Institute Brain Atlas, which has been at the forefront of brain sectioning and 3D reconstruction (Lein et al., 2007). Creating a reference framework and mapping anatomical positions of neurons and molecular phenotypes is an ongoing and very active field in brain research, and its extension to the cardiac brain -ICN - benefits from it. Our contribution also resembles reports that provide a data scaffold to enable adding new studies, such as Hsu and Bhandawat, 2016).

We find the rat ICN to be a bounded, relatively compact population within the context of the larger 3D anatomy of the heart. It is present on the hilum on the base of the heart, on both the left and right sides, as well as in the interatrial septum. The majority of neurons are on the posterior or dorsal surface of both atria, but more prominently on the left side, extending to the coronary sulcus separating the atria from the ventricles. Within the ICN, neurons tend to cluster and to some extent appear to be continuously distributed in the superior-inferior direction. Quantitative analysis of neuronal packing density suggests several distinct anatomical groupings that are conserved across individuals, but sex specific. The molecular phenotypes of these individual neurons are variable and diverse and reveal many previously unknown neuromodulatory regulators to be present. Spatial analysis suggests that many of these genes have specific gradients of expression, some of which is specifically associated with the neuronal packing density clusters, suggesting functional specificity by molecular phenotype and location within ICN.

### Anatomical mapping of intrinsic cardiac neurons in the 3D reconstructed whole heart

While qualitative and gross anatomical descriptions of the anatomy of the ICN have been presented, we here bring forth, at a cellular level, the first comprehensive atlas, and 3D reconstructed of the ICN in the rat. This work represents a quantum leap in a long history of attempts to understand the anatomical substrate upon which the neuronal control of cardiac function is built. There has not yet been an integrative effort to generate a comprehensive digitized neurocardiac anatomical atlas for any species, nor any effort to generate a histological foundation for molecular as well as functional mapping at the single neuron level for the whole heart of any species.

Prior studies have attempted to map the intrinsic cardiac ganglia, the autonomic afferent and efferent nerve innervation of these cardiac ganglia and cardiac myocardium. Commonly, immunohistochemical or immunofluorescent staining of a neuronal marker (acetylcholinesterase, tyrosine hydroxylase, choline acetyltransferase, PGP9.5, CGRP, SP) and a fluorescent tracer were used to examine sections, whole mounts or whole hearts with microscopic or macroscopic organ imaging. Mapping efforts of ICN and cardiac nerve innervation in small animals have included: mouse (Ai et al. 2007b, Hord et al 2008, Li et al., 2010, 2014; Rysevaite et al., 2011a, b), rat (Ai et al., 2007a, b; Cheng et al., 1999, 2004; Cheng and Powley, 2000, Pauza et al. 2002, Richardson et al., 2003), guinea pig (Pauza et al 2002, Hardwick et al., 2014; Steele et al., 1994), and rabbit (Saburkina et al., 2014,Pauziene 2016). Mapping of ICN was also performed in the hearts of larger animals and humans: dogs (Cardinal et al 2008, Pauza et al. 2002a,b,Singh et al. 2013, Xi et al. 1991), pigs (Arora et al 2003, Pauza et al. 2014, Nakamura et al. 2016), and humans (Pauza et al., 2000; Petraitiene et al., 2014; Singh et al., 1996, 2013). The ICN in all these species is characterized and summarized by Wake and Brack (2016). With these efforts, either restricted anatomical regions in tissue sections or flat whole-mounts were mapped at a microscopic level, or large gross anatomical regions that were mapped macroscopically.

While these studies demonstrate the location of the ICN, the extrinsic cardiac innervation by afferent and autonomic nerves, and the intrinsic cardiac innervation, they cannot however provide an anatomical substrate for integrating functional and molecular data with anatomically identified ICN neurons in their precise locations in the whole heart. In a sharp contrast, our work <u>*for the first time*</u> provides a novel 3D model to precisely integrate anatomical, functional and molecular data in the 3D digitally reconstructed whole heart with high resolution at the µm scale.

Recent technological advances in tissue clearing techniques make the heart transparent (Chung et al., 2013). Following tissue clearing, IHC and various microscopy techniques can be applied to map ICN and extrinsic and intrinsic cardiac nerves. Confocal and 2-photon imaging requires laser scanning, which is slow and hence practical for only small regions. ICN and extrinsic and intrinsic cardiac nerves may be beautifully imaged with light sheet microscopy, as demonstrated on the murine heart (Ding et al., 2018). This approach can present an ability to create an unbiased view of the myocardium at the single-cell level. However, there appears to be a trade-off between the physical width of the lightsheet (which can determine the axial resolution) and the size of the volume that can be imaged (Richardson and Lichtman, 2015). Hence, lightsheet microscopy with the highest resolution is limited to tissues that are a few hundred micrometers thick. In addition, the electrophoretic process in tissue clearing is not optimal for preservation of charged molecules, including DNA and RNA. Thus, our approach provides a much more comprehensive, reliable, and precise model to integrate anatomical, functional and molecular data.

### Molecular heterogeneity of intrinsic cardiac neurons: functional significance

The current understanding of ICN is that they modulate cardiac physiological functions of chronotropy, dromotropy, inotropy, and lusitropy. Several anatomical studies have hinted at the complexity of cardiac ganglia organization both in the cell body locations (Hasan, 2013; Hoover et al., 2009; Li et al., 2014; Pauziene et al., 2016; Rysevaite et al., 2011b, 2011a; Saburkina et al., 2014) and the complexity of afferent and efferent projections to and from the ICN (Cheng et al., 2004, 1999; Cheng and Powley, 2000; Lin et al., 2014; Li et al., 2014, 2010). While some work has hinted at the vast heterogeneity of the ICN from a functional perspective (Beaumont et al., 2013), there has not yet been a systematic assay capable of analyzing the molecular substrates that underlies this functional heterogeneity. For example, from a population perspective, it is exactly this type of heterogeneity that may determine the differences between patients who respond to vagal stimulation and those who do not. Furthermore, determination of the deficits in the ICN that drive cardiovascular pathology can lead to new avenues of therapy not only for patients with heart failure, but also with an eye towards preventative approaches, as the transition from health to disease is better understood from the perspective of the ICN (Herring, 2015;; Longpré et al., 2014). This work represents the first that we are aware of that attempts to elucidate the transcriptional heterogeneity of anatomically positioned ICN in the rat heart, providing anatomical and molecular substrates for functional heterogeneity of cardiac ICNs.

The distinct molecular phenotypes that are defined along the base-to-apex axis have not been previously described. That such an organizational pattern exists may be a result of the embryological development of the intrinsic cardiac ganglia, predominantly from the migration of neural crest cells that differentiate into neurons. The migrating neural crest cells enter the heart at the base, and spread out within the atria toward the apex (Fukiishi and Morriss-Kay, 1992; Hildreth et al., 2008). As in the brain, neurons in the heart likely follow chemical gradients to determine the direction and extent of migration, offering a possible explanation for why molecular phenotypes can be defined along the Z axis and opening the door for further work to determine which molecules make up these gradients. That cardiac conduction runs generally from base to apex may also suggest a functional implication for the separation of neurons in this fashion. In mouse hearts, not all intrinsic cardiac ganglia originate from migrating neural crest cells, especially those that influence nodal pacemakers (Hildreth et al., 2008). Further development of more comprehensive profiling of single neurons within their anatomical context will provide important information as to the role of both neural crest and non-neural crest neurons in the heart.

### Data analysis and integration framework

In order to piece together the relationship between 3D histological reconstructions and single cell molecular data, data storage and annotation must be carefully considered. Not only does the use of an anatomical ontology provide a means by which neurons can be anatomically referred to, but also makes it so that data from different hearts can be more readily compared. The generation of such data storage and annotation is not a trivial matter and requires collaboration between technicians, researchers, software developers, and data storage specialists. While purely mathematical and statistical methods for interpreting these data are crucial, the complexity quickly gets to a point where such approaches are challenging, making the ability to qualitatively interpret the data essential. To this end, MBF Bioscience has designed custom software, TissueMaker and Tissue Mapper, allowing for visualization of molecular and anatomical patterns. All processed and raw data are available via the SPARC Data Portal (https:\\data.sparc.science). The data structure adheres to the SPARC data format, and is curated by the SPARC data curation team to align the data to community ontologies, to ensure data integrity and to extract the Minimal Information Standard metadata.

The standardized data will enable mapping onto a 3D scaffold via a common coordinate framework in order to be used as a reference atlas for comparison across animals as well as across species and experimental conditions. These efforts are presently ongoing in coordination with the MAPCORE group of the SPARC program.

### Future directions for scalability and extension

We have established approaches that will now support scaling to acquire several male and female ICN providing a 3D framework as a foundational data resource - useful to developing detailed anatomical neural circuit/connectomic maps for neural control of the heart, and showing 3D distribution/gradients of molecular phenotypes. This unprecedented granularity in the understanding of cardiac neurons has the potential to unlock a new generation of cardiac therapeutics to treat or prevent all manner of cardiac pathology. To achieve the desired integration of different data types, the data from other approaches are visualized within the ICN of the 3D framework. The demonstration of cross-species application of the approach supports scaling to organisms with larger and more complex hearts such as pig and human, and to extension to other organs.

## METHODS

Figure 1 illustrates our two approaches as two graphical workflows. We established two multi-component pipelines to map neurons mapped from histological heart tissue sections. One approach optimizes precision of 3D heart shape and tissue section alignment for establishing a 3D reference framework. The second trades resolution of cardiac structure for, in addition to mapping neuron positions, the acquisition of projections/connectomic data and/or acquisition of neuron samples for molecular profiling. Figure 1A **shows the first pipeline** that acquires sections and images by using the novel Knife Edge Scanning Microscopy system (KESM, 3Scan) which maintains precision and eliminates most artifacts greatly improving anatomical rigor and reproducibility. Figure 1B **shows the second pipeline**, using cryostat sectioning, used here for a male and female rat heart. Images of these sections, including cell positions, can be stacked to create a whole heart volume with data that can be brought into the 3D reference systems created by the first approach.

### Knife Edge Scanning Microscopy

Images taken by KESM have great advantages for precision of gross morphology while resting on cell-level histology, ideal for developing 3D frameworks to hold additional data types from other approaches. Using Tissue Mapper software (MBF Bioscience) to map the position of each neuron as illustrated in Figure 2 we develop a comprehensive mapping of the precise extent and distribution of the ICN in the 3D framework of the heart, as described below:

#### Sample Preparation

In brief, a normal male rat heart was obtained fresh and subsequently immersed in 4% paraformaldehyde prior to whole-mount diffusion staining with Cresyl Echt Violet stain (0.05g in 50ml dH2O + 150µl glacial acetic acid, for 7 days) to enable visualization of the intrinsic cardiac neurons and ganglia.The tissue sample was subsequently paraffin embedded and KESM digitized.

#### Image Acquisition

The FFPE block was KESM digitized at slice thickness of 5 µm per z-slice. The FFPE block was mounted to a nano-precision XYZ robotic stage. The robotic stage moves samples along the XY axis towards a diamond knife ultramicrotome, which is coupled to a fiber optic cable. Hence, the cutting blade also serves as a source of illumination. There is custom-built objective, with a 5 mm field of view, and a tube lens that equates to a magnification of approximately 10x trained on the bevel of the diamond knife. The end of the tube lens is coupled to a CMOS TDI line scan color sensor with a 16K pixel resolution RGB output and a 5 µm x 5 µm pixel size.

The 5mm blade moves across the surface of the FFPE block, slicing and scanning simultaneously, capturing one continuous line (or strip) of image data at a time, to generate an image tile comprised of 10,000 pixels, with each pixel representing 0.5 μm. After the strip has been fully sectioned and imaged, the stage is moved to position the adjacent region of the heart in line for sectioning and imaging. This process is repeated until the entire heart has been sectioned.

#### Image Processing

Post-processing of the data utilizes the precise spatial alignment provided by the KESM technique to generate 2D image planes, which were subjected to denoising and artefact reduction. Individual image planes were assembled together into a 3D volume, enabling quantification of morphological details over large anatomical distances. To achieve this, individual KESM image tiles acquired from each XY location at each Z position were automatically aligned and stitched into 2D image planes, cropped to remove excess image data that did not contain the heart, and then assembled into a 3D image volume with 35:1 JPEG2000 compression (Biolucida Converter, MBF Bioscience, Williston). These image volumes were annotated using Tissue Mapper as described below.

### Software Development and Neuron Mapping

A custom suite of computational mapping programs have been developed for mapping neurons (or any cell of interest) in organs including the heart: TissueMaker and Tissue Mapper. These were, and continue to be, evolved from the tools MBF Bioscience has developed for brain mapping, such as Neurolucida and BrainMaker.

The rat heart sectioned at TJU as in Figure 1B was run through the TissueMaker and Tissue Mapper pipeline. By contrast, as in Figure 1A the KESM image data from 3Scan did not require the alignment step in TissueMaker because it was already spatially aligned (see imaging description above).

Using Tissue Mapper software, precise locations of each cardiac neuron were mapped in all sections in which neurons are present. In addition to marking the cell location, numerous regions selected from the comprehensive ontology first generated for the Cardiac Physiome Project (Hunter and Smith, 2016) were mapped. Of the thousands of ontological features that were available, less than 40 were selected for these initial representations in order to simplify the images and to test the pipeline more efficiently. On each section, researchers traced key features (aorta, pulmonary vessels, atrial borders, etc.) and identified neurons based upon a combination of Nissl staining and morphology. The TissueMaker and Tissue Mapper software are then able to generate 3D wireframes of the hearts with neurons positioned in context. Attributes like color and shape can be customized in these reconstructions and quantitative spatial data can be obtained. By mapping anatomical fiducial information alongside the neuron locations, the extent and location of the neurons within the larger context of the entire heart could be viewed as a 3D representation, as seen in the Figures for the present work.

### Cryosectioning and Embedding

In our initial efforts with Heart A (male) and Heart B (female) we learned that, unlike the brain, the distortions of the heart are far less homogenous and symmetric, which can interfere with data visualization and comparison between specimens. Thus we developed the method described below to keep the chambers inflated. We also discovered that the embedding media needs color added to permit image segmentation. Optimal Cutting Temperature media (OCT, TissueTek; VWR 25608-930) is added to an embedding mold, to cover the bottom of the mold and kept on dry ice. Three concentrations of OCT diluted in 1x PBS are prepared 25% 50%, and 100%. The 100% OCT preparation should include a few drops of green food grade dye to permit optimal image segmentation of any subsequent blockface images. The animal is sacrificed using rapid decapitation after 60 seconds of exposure to 5% isoflurane. The heart is immediately excised and submerged in room temperature 1x PBS for 30 seconds or until the majority of blood is pumped out of the chambers. The still beating heart is transferred to 25% OCT, and lightly agitated for 30-60 seconds. This is then transferred to a 50% OCT solution, and lightly agitated for 30-45 seconds. Ideally, the heart should still be beating at this point. The large chambers of the heart were injected with colored OCT using a 14-16 gauge blunt needle on a syringe in order to mitigate structural collapse during cryosectioning. Next, the heart is placed in a chilled embedding mold. Room-temperature OCT is added to completely submerge the heart. The mold is then placed in a slurry of dry ice and methanol to promote rapid freezing. Care was taken to avoid allowing methanol to come into contact with the OCT in the block as this will compromise the structural integrity of the OCT once frozen. Note that holding the block near liquid nitrogen, but not submerging it, is also a means of rapid cooling. After the OCT is completely frozen, it is covered with aluminum foil and then in a plastic wrap to prevent accumulation of condensation in the mold. The mold is transferred to a −80°C freezer. Ideally the entire process should happen within 5-10 minutes to mitigate RNA degradation, which will be necessary for future investigations of single cell transcriptomics using laser capture microdissection.

### Slide Preparation and Image Processing

Heart A was sectioned from base to apex at 20µm, yielding nearly 800 sections, with corresponding blockface images, and mounted onto 400 slides (two sections per slide). In order to appreciate a finer level of detail and clarify tissue staining, Heart B was sectioned at 10 um yielding nearly 1600 sections mounted on 800 slides with blockface images for each section.

Each slide was stained with 0.1% cresyl violet and dehydrated using increasing concentrations of ethanol and xylene. Cover slips were added using mounting media and slides were then imaged using a slide scanner equipped with 20x Olympus objective (N.A.= 0.75; Bliss-200, MBF Bioscience, Williston, VT). On average, 2,000 image tiles were automatically acquired and stitched to create a high-resolution whole slide image containing two heart sections per image. Using TissueMaker software, the section images were then extracted from the whole slide image, cropped to a uniform size, the perimeter automatically segmented, and aligned spatially by identifying the centroid of each section to generate an image stack. The heart image volume was then shared with TJU and UCF for further mapping and segmentation using custom-developed software (Tissue Mapper; MBF Bioscience). The software application includes annotation tools for automatically or manually drawing regions, placing markers to indicate cell positions and other discrete points, and the ability to import comprehensive lists of regions as a text or comma-delimited file or via direct integration with the SciCrunch database (https://scicrunch.org/).

### Laser Capture Microdissection of Single Neurons

In order to isolate neurons while maintaining their anatomical origin in three dimensional space, it was necessary to use laser capture microdissection (Arcturus, ThermoFisher). Neurons were visualized using a rapid cresyl violet stain that highlighted the histological appearance of neurons and maintained RNA quality. Single neurons were collected on Capsure HS caps and the cells lysed right on the cap within 15 minutes after laser capture using lysis buffer from the CellsDirect DNA extraction kit (Life Technologies).

### Transcriptional assay of laser captured single neurons using multiplex RT-qPCR

RNA-seq performed on laser-captured single cells has only recently been demonstrated and was not available during these experiments (Foley et al. 2019). In order to assay single neurons from laser capture, it was necessary to apply multiplex RT-qPCR to ensure the sensitivity to detect several genes of interest given the small amount of RNA per sample. The Biomark microfluidic system (Fluidigm, San Francisco, CA) was used for all gene expression assays. After reverse transcription and whole transcriptome amplification (Qiagen, Hilden, Germany), the samples were processed through the Biomark system following manufacturer suggested protocols. Quality control of RT-qPCR results included filtering through melt-curve analysis along with automatic C_t_ thresholding to determine the limit of detection.

## Supporting information

List of Supplemental Files

## ACKNOWLEDGEMENTS

We would like to thank Dr. Mahyar Osanlouy of the Auckland Bioengineering Institute for ongoing efforts in the process of fitting segmented data to the heart scaffold. Financial support for this work was provided by the National Institutes of Health under the Stimulating Peripheral Activity to Relieve Conditions (SPARC) program. Grant OT2 OD023848. NHLBI Grant U01HL133360 to JSS and RV. The funders had no role in study design, data collection and interpretation, or the decision to submit the work for publication.

