## Supplementary material for "A Comprehensive Integrated Anatomical And Molecular Atlas Of Rodent Intrinsic Cardiac Nervous System": List of Supplemental Files

**Supplemental Video 1:** Rotating model of the Rat ICN in cardiac context. Cardiac features have been contoured and colored accordingly, with mapped neurons marked in yellow. Partial projection of the neurons at the base of the heart are also shown.

**Supplemental Video 2:** Identification of Neuronal Clusters in the Heart. Visualization of neuronal clusters based on packing density by use of partitioning around medoids (PAM) algorithm. Clusters are shown as a flat map projection as well as their mapped locations within the ICN.

**Supplemental Video 3:** Visual representation of both mapped and sampled neurons in the rat ICN, including flat map projections. Neurons are colored based off of clusters identified through PAM algorithm as described.

**Supplemental Table 1:** File containing the raw Ct values for the sampled neurons.

**Supplemental Table 2:** File containg the normalized qPCR data (-ddct). This file also details the metadata such as the X,Y,Z coordinates, slide number, and cell count per sample.

**Supplemental Figure S1:** Microscopic images are available for all the collected samples. In the dataset on the SPARC Data Portal, within the "Samples" folder, there is a document called "Sample_Collection_Information" (Bottom Panel) and another folder titled "Supplementary Sample Acquisition Images". The top level of this folder is shown on the top panel (left), where the folder names correspond to the slide number of the sample in question (top panel, left and bottom panel, column A). Within that folder is a folder for each slide region (top panel, top right, and bottom panel, column B). Within each folder for a slide region is a folder for each sample containing the images taken for the acquisition of that sample (top panel, bottom right, and bottom panel, column C). For each sample, the corresponding folder contains images showing the area of interest, the neurons marked before they are collected, the neuron after it was collected on the collection cap, the remaining tissue after the neurons had been picked, as well as images showing the tissue before and after an additional UV cut which aided in the mapping of samples later on (bottom panel columns F-K). All images are labeled with the date they were collected followed by a number that is listed in the spreadsheet to identify the images.
