## Supplementary figures and images for "A Comprehensive Integrated Anatomical And Molecular Atlas Of Rodent Intrinsic Cardiac Nervous System"

### Supplemental Figure 1

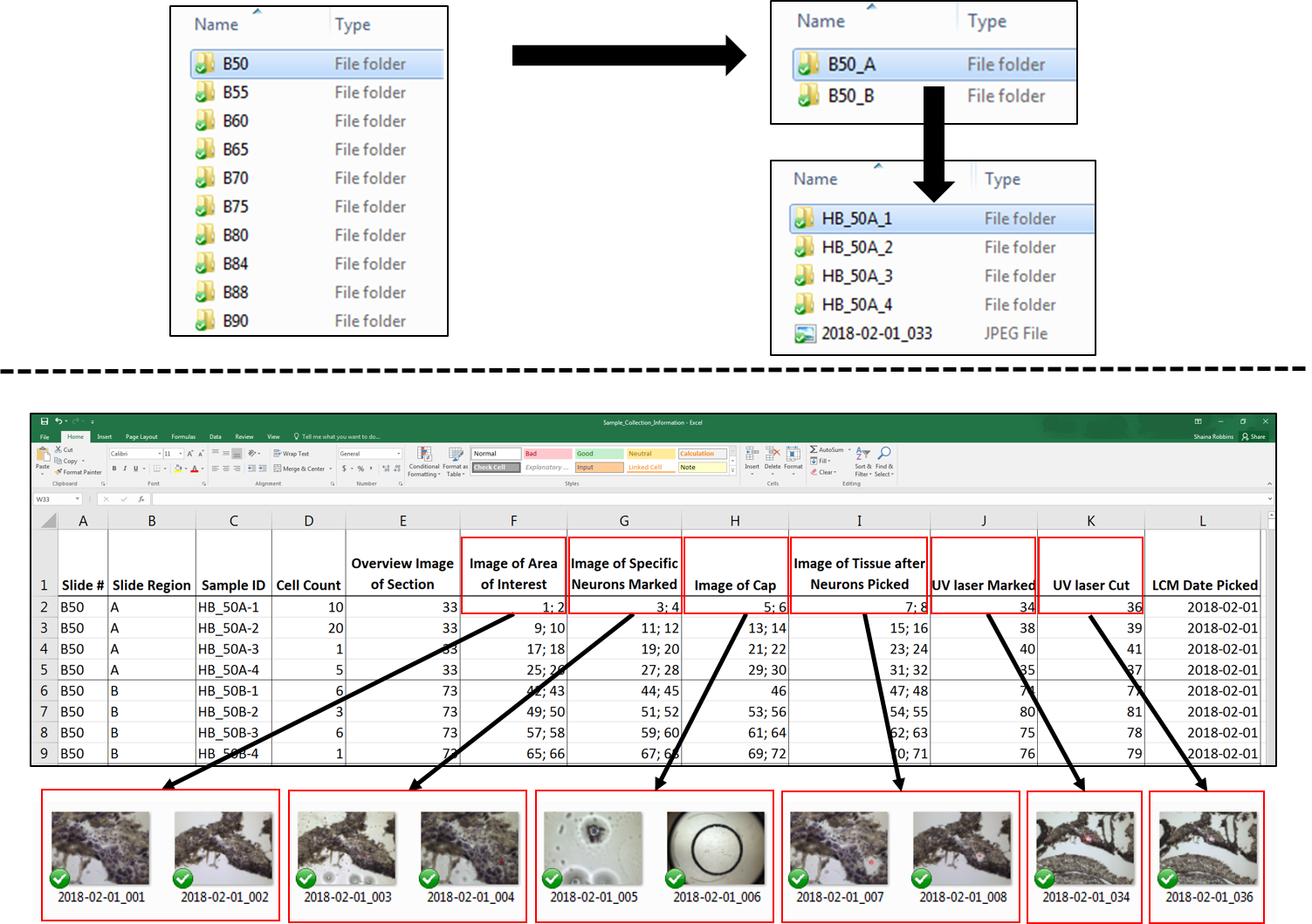
